# The transcriptome of rat hippocampal subfields

**DOI:** 10.1101/2021.06.23.449669

**Authors:** João P. D. Machado, Maria C.P. Athie, Alexandre H. B. Matos, Iscia Lopes-Cendes, André. S. Vieira

## Abstract

The hippocampus comprises several neuronal populations such as CA1, CA2, CA3, and the dentate gyrus (DG), which present different neuronal origins, morphologies, and molecular mechanisms. Laser capture microdissection (LCM) allows selectively collecting samples from target regions and eliminating unwanted cells to obtain more specific results. LCM of hippocampus neuronal populations coupled with RNA-seq analysis has the potential to allow the exploration of the molecular machinery unique to each of these subfields. Previous RNA-seq investigation has already provided a molecular blueprint of the hippocampus, however, there is no RNA-seq data specific for each of the rat hippocampal regions. Serial tissue sections covering the hippocampus were produced from frozen brains of adult male Wistar rats, and the hippocampal subfields CA1, CA2, CA3, and DG were identified and isolated by LCM. Total RNA was extracted from samples, and cDNA libraries were prepared and run on a HiSeq 2500 platform. Reads were aligned using STAR, and the DESeq2 statistics package was used to estimate gene expression. We found evident segregation of the transcriptomic profile from different regions of the hippocampus and the expression of known, as well as novel, specific marker genes for each region. Gene ontology enrichment analysis of CA1 subfield indicates an enrichment of actin regulation and postsynaptic membrane AMPA receptors genes indispensable for long-term potentiation. CA2 and CA3 transcripts were found associated with the increased metabolic processes. DG expression was enriched for ribosome and spliceosome, both required for protein synthesis and maintenance of cell life. The present findings contribute to a deeper understanding of the differences in the molecular machinery expressed by the rat hippocampal neuronal populations, further exploring underlying mechanisms responsible for each subflied specific functions.

## 1 INTRODUCTION

The Hippocampus is one of the most studied structures of the nervous system. It is involved in diverse functions such as spatial navigation, processing of memories and emotional responses, and presents subregions or subfields arranged in a complex circuit (Witter et al., 2014). Each of these subfields, CA1, CA2, CA3, and the dentate gyrus (DG), have different neuronal origins, morphologies, and molecular mechanisms (Hayashi et al., 2015; *Microcircuits of the Hippocampus,* 2016). CA1, CA2 and CA3 subfields are mainly composed of pyramidal cells, creating an output circuitry for the hippocampus. On the other hand, DG cells are the input sub-region, consisting mainly of granule cells *(Microcircuits of the Hippocampus,* 2016). All these cells are in constant communication with different types of inhibitory neurons, which control the excitatory waves produced on the hippocampus (Albrecht et al., 2020; Booker & Vida, 2019). Therefore, an accurate selection of cell populations would preserve regional characteristics, potentially helping to better understand differences in the hippocampal functionality.

Laser capture microdissection (LCM) has enabled an accurate microstructure isolation using a laser coupled to a microscope, which cuts accordingly to a trajectory predefined by the user (Espina et al., 2006). LCM technique allows selectively collecting samples from target regions and eliminating unwanted cells to obtain more specific results (Datta et al., 2015). The LCM process does not alter the integrity of a collected sample, thus it is an excellent method to collect cells or cell subfields preserving RNA integrity (Espina et al., 2006). The use of regional LCM to isolate CA1, CA2 and CA3 pyramidal layers, and DG granular layer would allow the exploration of the molecular machinery unique to each of these subfields. In conjunction with transcriptomic tools, it is possible to define expression characteristics of a given neuronal population and to obtain region-specific transcriptomes, essential for a deeper insight into hippocampus function.

Despite all hippocampal transcriptome investigations done so far, most have not explored the regional heterogeneities of this structure. Furthermore, gene expression features such as gene ontology and quantitative analysis of transcripts remain unclear and need further investigation. Studies separating the hippocampal subfields have been essentially performed by microarrays (Greene et al., 2009; Lein et al., 2004; Nakamura et al., 2011; X. Zhao et al., 2001), which have limitations when compared to RNA-seq experiments (Rampil, 2007). Previous RNA-seq investigation has already provided a molecular blueprint of the mouse hippocampus (Cembrowski et al., 2016), however, there is no RNA-seq data specific for the rat hippocampal subfields.

Here, we explore the gene expression heterogeneity of rat hippocampus by using LCM to isolate its different subfields (CA1, CA2, CA3, DG), and by tracing a profile of those regions transcriptome. We also provide insights into the transcriptional organization of specific enriched ontologies or pathways for each subfield and interpret possible functional characteristics associated with such expression patterns.

## 2 METHODS

### 2.1 Animals and Laser microdissection (LCM)

In this study, three month old male Wistar rats were housed in a ventilated environment (12□h/12□h light cycle) with *ad libitum* access to standard rodent chow and water. All procedures were executed according to the ethical standards for animal experimentation at the University of Campinas-UNICAMP (Brazilian federal law 11.794 (10/08/2008 – Animal Use Ethics Committee protocol 2903-1). Four rats (n=4) were anesthetized with isoflurane (2% isoflurane, 98% oxygen at 1 liter/min) and decapitated using a small animal guillotine. Then, brains were snap-frozen at −55°C and posteriorly processed in a cryostat (Leica Biosystems – Wetzlar, Germany) to obtain 60-μm serial sections covering the entire hippocampus.

For laser microdissection, tissue sections were collected in PEN membrane-covered slides (Life Technologies^®^, Thermo Fisher Scientific – Waltham, USA), stained with Cresyl Violet, dehydrated with ethanol, and stored at −80□°C. The hippocampal subfields were identified according to Paxinos Rat Brain Atlas 7th edition (Paxinos & Watson, 2013) and delimited with a Palm (Zeiss^®^ – Jena, Germany) system. We followed the hippocampal microdissection methods outlined by (Vieira et al., 2016), concerning the hippocampal subfields. Finally, tissue was mechanically collected in separate microcentrifuge tubes using a surgical microscope and micro-forceps. We collected the granular subfield of the DG and the pyramidal subfields of CA1, CA2 and CA3.

### 2.2 Library preparation and Next-generation sequencing

RNA was extracted with TRIzol (Thermo Fisher Scientific – USA) using the manufacturer instructions, then cDNA libraries were reverse-transcribed from 200□ng of extracted RNA using TruSeq Stranded Total RNA LT (Illumina^®^ – USA), according to the manufacturer instructions. Each sample library was uniquely barcoded to be sequenced in a HiSeq^®^ 2500 (Illumina^®^) in High Output mode, producing 100-bp paired-end sequences.

### 2.3 Data processing and differentially expressed genes (DEGs)

All sequenced reads were aligned using the StarAligner 2.6 program (https://github.com/alexdobin/STAR, RRID:SCR_004463) with the Rattus norvegicus genome 3.1 (Rnor6 Ensembl release 6.0 – https://www.ensembl.org/Rattus_norvegicus/lnfo/lndex). Subsequently, DESeq2 package version 3.12 (http://bioconductor.org/packages/release/bioc/html/DESeq2.html, RRID:SCR_015687) was used to calculate differentially expressed genes (DEGs) between subfields and carry statistical analysis. DEGs were submitted to enrichment analysis using clusterprofiler package (https://bioconductor.org/packages/release/bioc/html/clusterProfiler.html, RRID:SCR_016884) and the functional profiles were classified into KEGG Pathways and GO Terms – Biological Processes (BP). Differential gene expression was considered significa when adjusted p-value<0.05. For all enrichment analyses, terms and pathways were considered significantly different when p<0.05 (after adjustment for multiple comparisons).

## 3 RESULTS

### 3.1 The Sequencing Run

A total of 253,341,913 100-bp paired-end reads were produced from all samples, averaging 15.7 million paired-end reads per sample. The average sequence alignment was 77,5% and read counts per gene were used to estimate gene expression and statistical analysis in DESEQ2.

### 3.2 Identification of DEGs and Samples Visualization

We ran six pairwise comparisons in DESEQ2, comparing all subfields between themselves, and obtained the following DEGs results (adjusted p<0,05) 2863 (CA1xCA2), 4318 (CA1xCA3), 1847 (CA2xCA3), 5361 (CA1xDG), 4815 (CA2xDG) and 7120(CA3xDG). For a complete list of all DEGs refer to Supplementary Table 1. The quantities of DEGs, upregulated and downregulated, are represented in Figure (1A). All uniquely and commonly DEGs are represented in the Venn diagram (Figure 1B). PCA displays an evident clustering of samples of the different regions of the hippocampus: CA1, CA2, CA3, and DG (Figure 1C).

**Figure 1.**
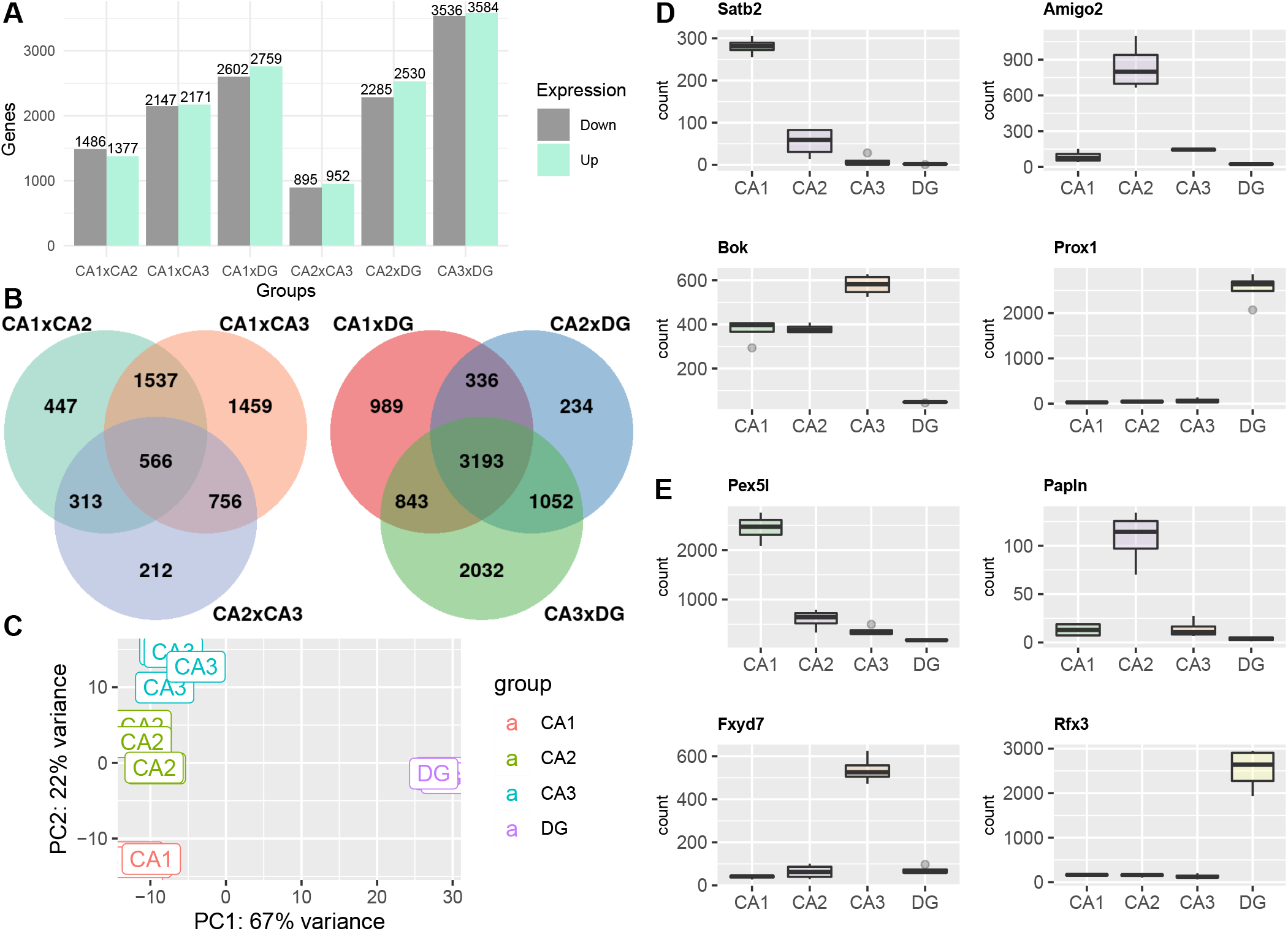
Intragroup and intergroup variability. **(A)** Barplot of the transcriptomics data displaying the number of differentially expressed transcripts identified in each hippocampal subfield comparison. **(B)** Venn diagram representing common and unique differentially expressed genes in all comparisons. **(C)** PCA graphic for hippocampal gene expression data displaying an evident cluster of samples on the different regions of the hippocampus. **(D)** Plotcounts of distinct marker genes for each hippocampal subfield.

### 3.3 Molecular markers for hippocampal subfields

Known molecular markers such as *‘Satb2’, ‘Amigo2’, ‘Bok’* and *‘Prox1’*, for subfields CA1, CA2, CA3, and DG respectively (Hamilton et al., 2017), presented unique expression for those subfields in our dataset (Figure 1D). In addition, we evaluated potential new marker genes among the DEGs in the dataset (>4 fold change comparing a subfield against all others) unique to each subfield (Figure 1E). For the whole set of potential marker genes, please refer to Supplementary Table 2.

### 3.4 Pyramidal to pyramidal subfiled comparisons

#### CA1xCA2

GO analysis revealed a distribution of 111 significant (p.adjust < 0.05) BPs for CA1xCA2 considering all DEGs (See Supplementary Table 3). Top hits were *‘regulation of membrane potential’, ‘synapse organization’, ‘axonogenesis’,* and other noteworthy BPs like *‘neuronal death’* and *‘neurotransmitter secretion’*. Splitting DEGs into downregulated or upregulated genes lists and using CA1 as reference evidenced other enriched terms. The downregulated GO networks association revealed core BP terms like: *‘synaptic vesicle cycle’, ‘ATP metabolic process’, ‘generation of precursor metabolites and energy^1^, ‘regulation of neurotransmitter levels’* and *‘regulation of membrane potential*’ (Figure 2A). All these functions and genes are activated in CA2 but downregulated in the CA1 gene set expression list. On the other hand, for CA1 upregulated genes, GO networks showed *‘cell junction assembly’, ‘regulation of membrane potential’*, and *‘synapse organization’* as enriched terms (Figure 2B). We also performed a KEGG pathway analysis separately on splitted gene lists and the list of top enriched pathways is demonstrated in Figures (2C and 2D), for a complete list of KEGG pathways refer to Supplementary Table 4. The downregulated genes in CA1 enriched pathways include *‘pathways of neurodegeneration’, ‘thermogenesis’, ‘rap1 signaling pathway’, ‘oxidative phosphorylation’, ‘carbon metabolism’, ‘synaptic vesicle cycle’,* and *‘glycolysis/gluconeogenesis’*. Meanwhile, *‘calcium signaling pathway’, ‘axon guidance’, ‘regulation of actin cytoskeleton’, ‘cAMP signaling pathway’, ‘glutamatergic synapse’, ‘long-term potentiation’, ‘adherens junction’,* and *‘glycosaminoglycan biosynthesis – heparan sulfate/heparirn’* appears upregulated for CA1.

**Figure 2.**
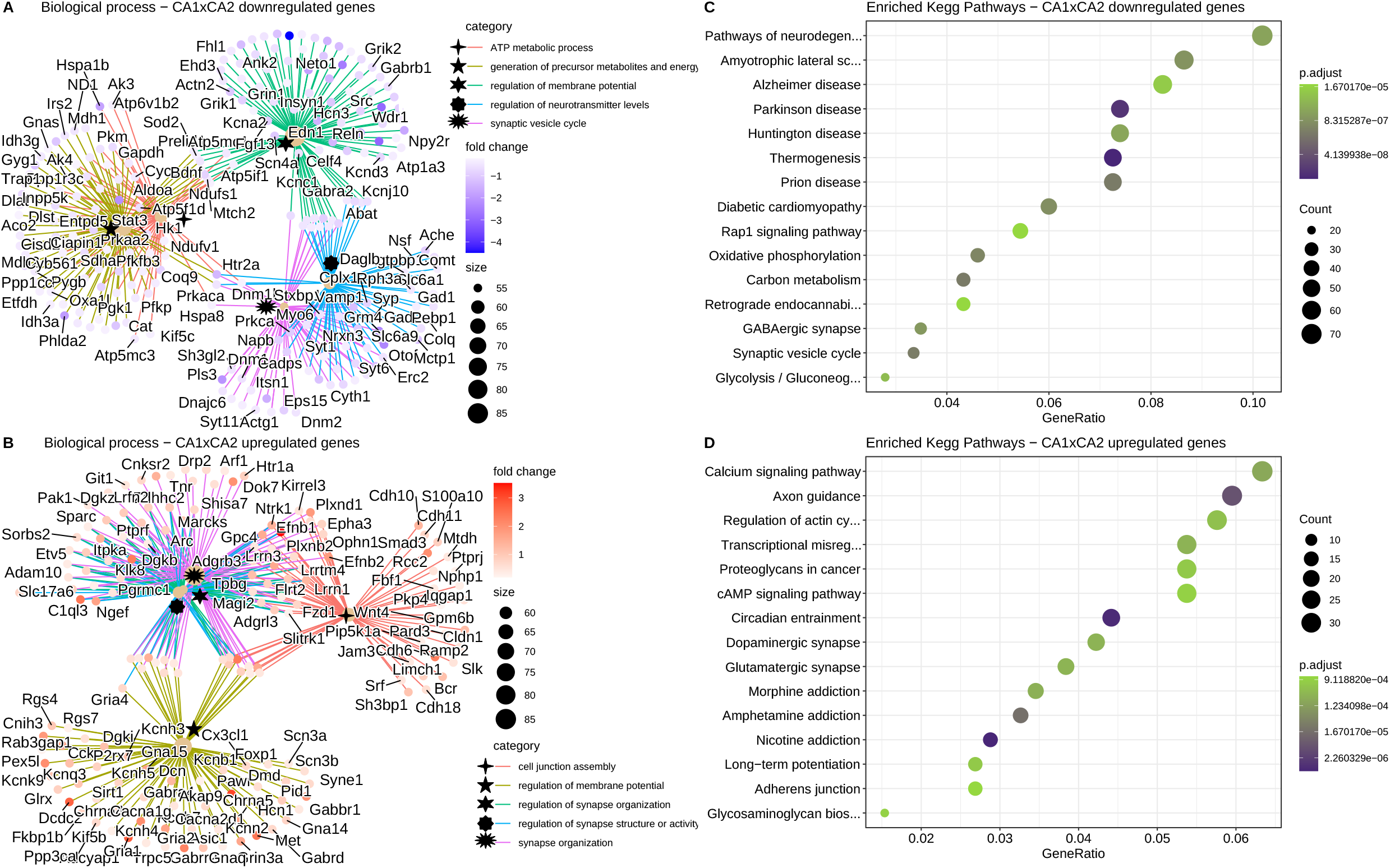
Top enriched biological process and KEGG pathway of CA1×CA2 differentially expressed genes. **(A)** A net plot of five enriched biological processes from GO analysis of upregulated genes(CA1xCA2). **(B)** A net plot of five enriched biological processes of downregulated genes(CA1xCA2) and their differentially expressed genes. **(C)** A dot plot of top enriched KEGG pathways in CA1xCA2 downregulated genes. **(D)** A dot plot of top enriched KEGG pathways in CA1xCA2 downregulated genes. All enriched pathways and biological processes displayed reached adj.p-value<.05. For the whole set of enriched biological processes and pathways, please refer to Supplementary Table 3 and 4.

#### CA1xCA3

We found 118 biological processes significantly enriched (p.adjust <0.05) for CA1xCA3’s GO analysis (See Supplementary Table 3). Among all the enriched functions, we highlight terms such as: *‘synapse organization’, ‘regulation of membrane potential’, ‘positive regulation of cell projection organization’,’exocytosis’* and *‘synaptic vesicle cycle’.* Downregulated genes list for CA1 provided enriched terms like: *‘ATP metabolic process’, ‘generation of precursor metabolites and energÿ, ‘oxidative phosphorylation’, ‘regulation of membrane potential’* and *‘synaptic vesicle cycle’* (Figure 3A). On the other hand, GO analysis for upregulated genes in CA1 displays the following enriched biological terms: *‘actin filament organization’, ‘positive regulation of cell projection organization’, ‘regulation of synapse organization’, ‘regulation of synapse structure or activity’* and *‘synapse organization’* (Figure 3A). In addition, we found enriched KEGG pathways for downregulated genes in CA1 such as: *‘pathways of neurodegeneration’, ‘thermogenesis’, ‘rap1 signaling pathway’, ‘oxidative phosphorylation’, ‘carbon metabolism’, ‘synaptic vesicle cycle’,* and *‘glycolysis/gluconeogenesis’* (Figure 3C). *‘axon guidance, ‘regulation of actin cytoskeleton’, ‘cAMP signaling pathway’, ‘glutamatergic synapse’, ‘cholinergic synapse’, ‘adherens junction’,* and *‘glycosaminoglycan biosynthesis – heparan sulfate/heparirin’* are the KEGG enriched pathways for upregulated genes in CA1 (Figure 3D). For a complete list of KEGG terms refer to Supplementary Table 4.

**Figure 3.**
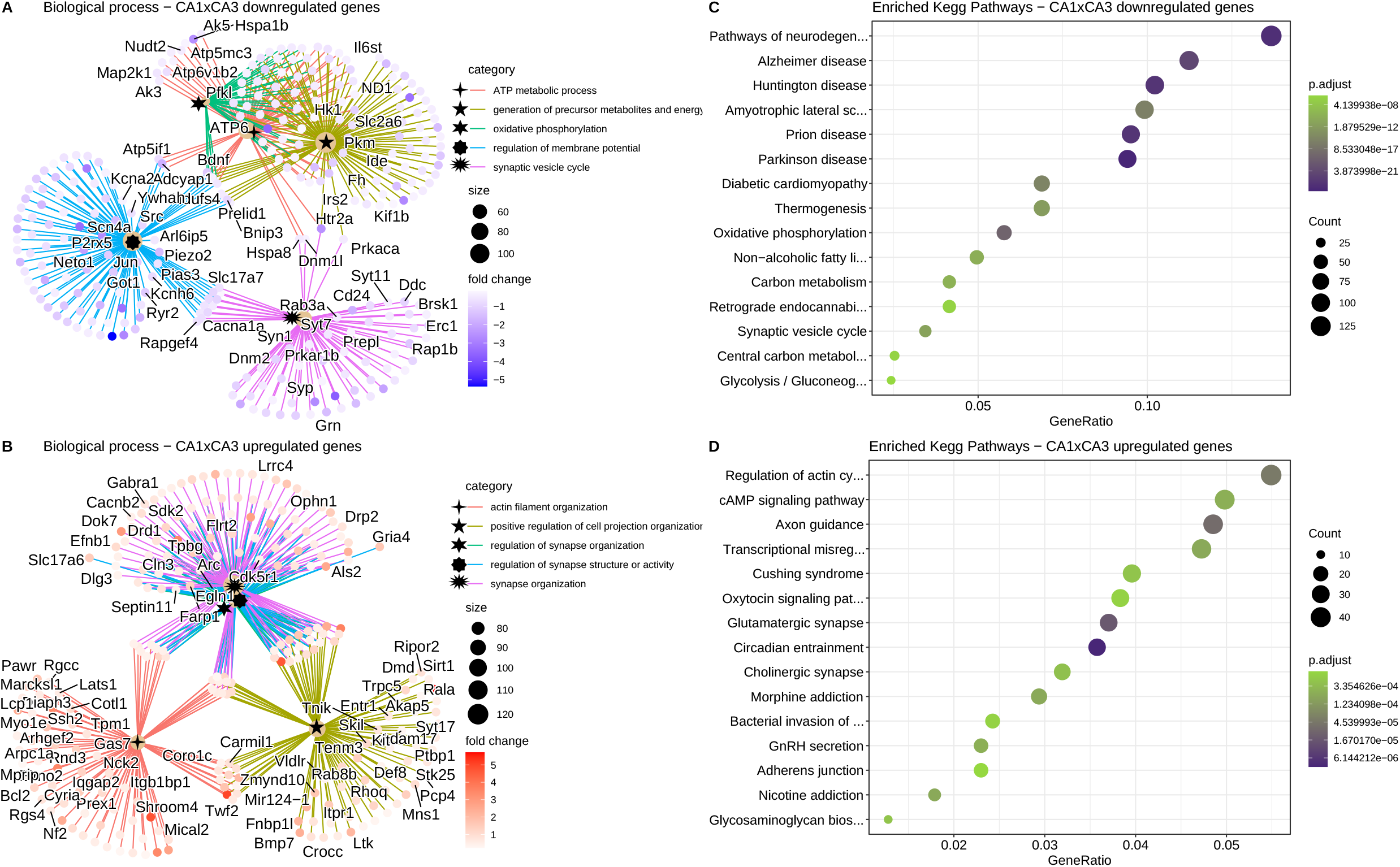
Top enriched biological process and KEGG pathway of CA1×CA3 differentially expressed genes. **(A)** A net plot of five enriched biological processes from GO analysis of upregulated genes(CA1xCA3). **(B)** A net plot of five enriched biological processes of downregulated genes(CA1xCA3) and their differentially expressed genes. **(C)** A dot plot of top enriched KEGG pathways in CA1xCA3 downregulated genes. **(D)** A dot plot of top enriched KEGG pathways in CA1xCA3 downregulated genes. All enriched pathways and biological processes displayed reached adj.p-value<.05. For the whole set of enriched biological process and pathways, please refer to Supplementary Table 3 and 4

#### CA2xCA3

A total of 110 biological processes significantly enriched (p.adjust <0.05) were found for CA2xCA3’s GO analysis (See Supplementary Table 3). The main BP involved for all DEGs of CA2xCA3 were as follows: ‘synapse organization’, *‘regulation of membrane potential’, ‘actin filament organization’, ‘regulation of GTPase activity’, ‘neurotransmitter transport’,* and *‘potassium ion transport’.* The downregulated genes for CA2 revealed GO terms associated with: *‘neuron projection guidance’, ‘synapse organization’,* and a core of three terms related to *‘negative regulations of cell projection and development’* (Figure 4A). On the other hand, we found upregulated genes in CA2 related to such BPs: *‘positive regulation of cell projection organization’, ‘dendrite development’, ‘synapse organization’,* and *‘actin filament organization’* (Figure 4B).

**Figure 4.**
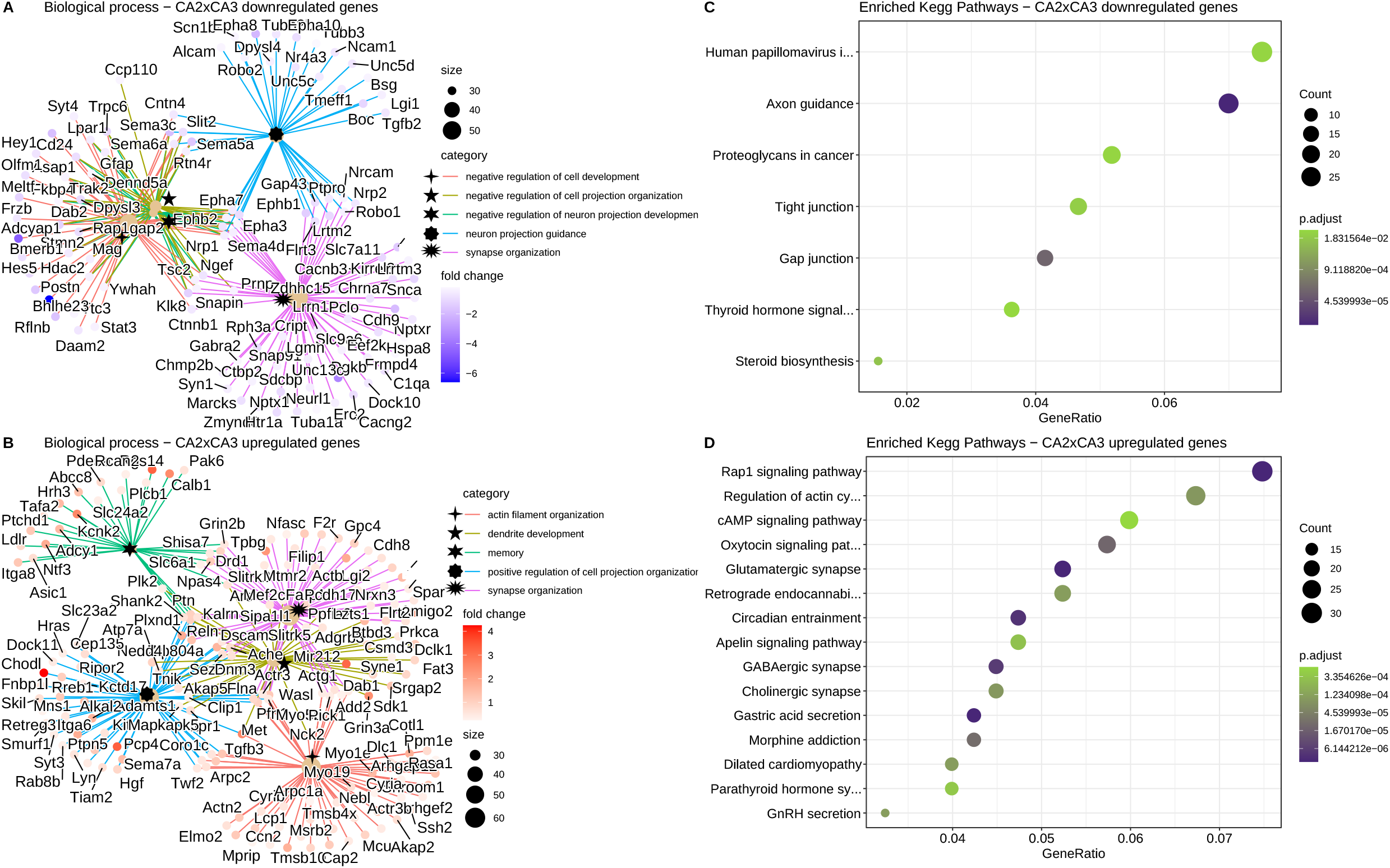
Top enriched biological process and KEGG pathway of CA2xCA3 differentially expressed genes. **(A)** A net plot of five enriched biological processes from GO analysis of upregulated genes(CA2xCA3). **(B)** A net plot of five enriched biological processes of downregulated genes(CA2xCA3) and their differentially expressed genes. **(C)** A dot plot of top enriched KEGG pathways in CA2xCA3 downregulated genes. **(D)** A dot plot of top enriched KEGG pathways in CA2xCA3 downregulated genes. All enriched pathways and biological processes displayed reached adj.p-value<.05. For the whole set of enriched biological process and pathways, please refer to Supplementary Table 3 and 4

Furthermore, top enriched KEGG pathways for downregulated genes in CA2 were: *‘axon guidance’, ‘proteoglycans in cancer’, ‘tight junction’, ‘gap junction’, ‘thyroid hormone signaling’,* and *‘steroid biosynthesis’* (Figure 4C). For upregulated genes in CA2, we found: *‘Rap1 signaling pathway’, ‘regulation of actin cytoskeleton’, ‘cAMP signaling pathway’, ‘retrograde endocannabinoid signaling’, ‘glutamatergic synapse’ ‘apelin signaling pathway’ ‘cholinergic synapse’,* and *‘GABAergic synapse’* (Figure 4D). For a complete list of KEGG terms refer to Supplementary Table 4.

### 3.5 Granular to pyramidal subfield comparisons

GO analysis revealed the following distribution: 128 significantly enriched BP (p.adjust <0.05) for CA1xDG; 133 significantly enriched BP (p.adjust <0.05) for CA2xDG; and 102 significantly enriched BP (p.adjust <0.05) for CA3xDG (See Supplementary Table 3). We also did the KEGG pathway analysis of solemnly up and solemnly down gene lists separately to obtain more specific terms about the differences between the subfields and to improve the search for terms (Figure 5). For a complete list of KEGG pathways refer to Supplementary Tables 4.

**Figure 5.**
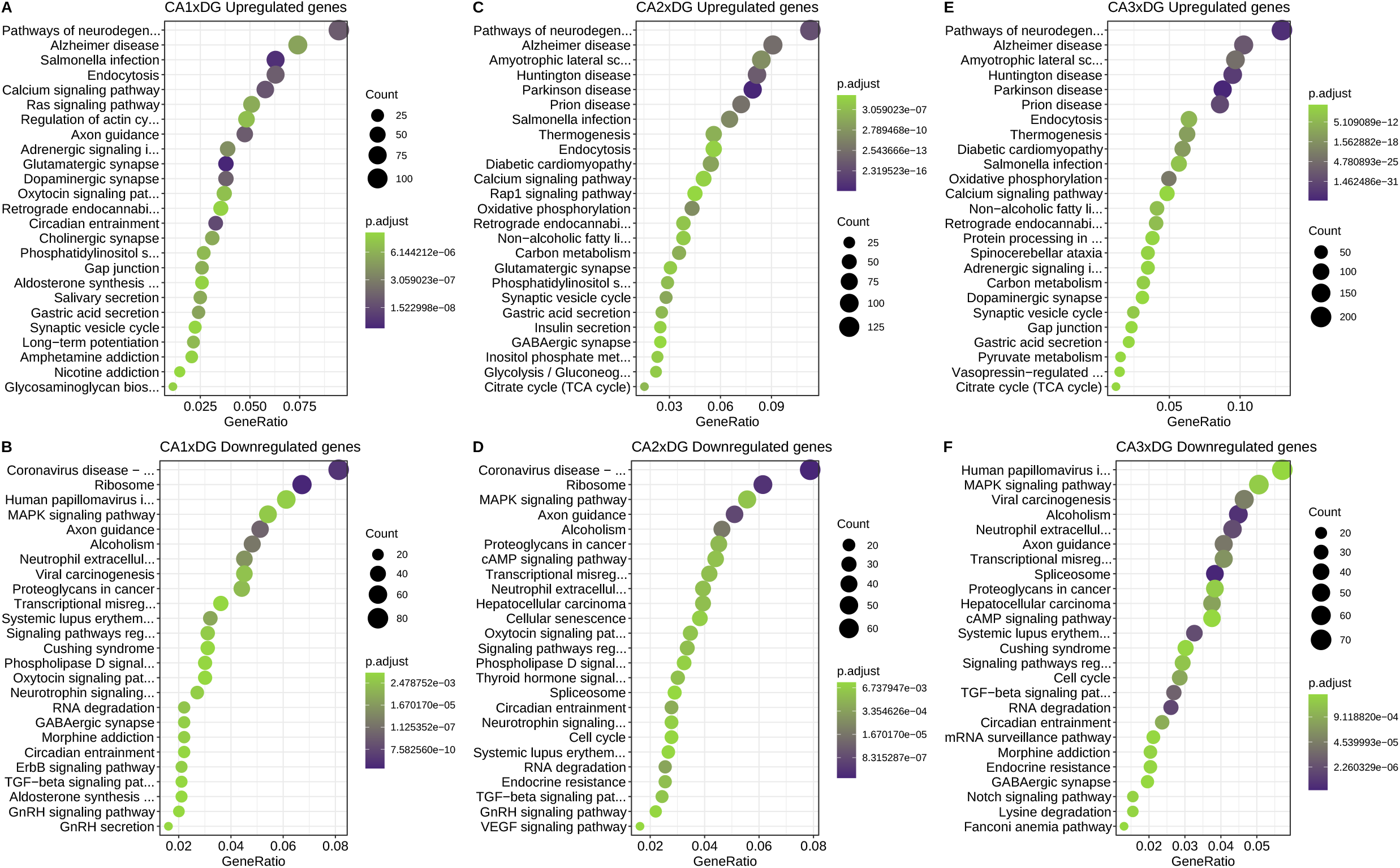
Enriched KEGG pathway of upregulated and downregulated genes in CA’sxDG comparisons. **(A)** A dot plot of top enriched KEGG pathways in CA1xDG upregulated genes. **(B)** A dot plot of top enriched KEGG pathways in CA1xDG downregulated genes. **(C)** A dot plot of top enriched KEGG pathways in CA2xDG upregulated genes. **(D)** A dot plot of top enriched KEGG pathways in CA2xDG downregulated genes. **(E)** A dot plot of top enriched KEGG pathways in CA3xDG upregulated genes. **(F)** A dot plot of top enriched KEGG pathways in CA3xDG downregulated genes. All enriched pathways and biological processes displayed reached adj.p-value<.05. For the whole set of enriched biological process and pathways, please refer to Supplementary Table 3 and 4

#### CA1xDG

We found the following KEGG pathways for positively regulated genes in CA1: *‘pathways of neurodegeneration’, ‘endocytosis’, ‘calcium signaling pathway’, ‘ras signaling pathway’, ‘regulation of actin cytoskeleton’, ‘axon guidance’, ‘glutamatergic synapse’, ‘dopaminergic synapse’, ‘cholinergic synapse’, ‘synaptic vesicle cycle’,* and *‘longterm potentiation’* (Figure 5A). Next, KEGG analysis for downregulated genes for CA1 reveals terms like: ‘ribosome’, ‘MAPK signaling pathway’, ‘axon guidance’, ‘phospholipase D signaling pathway’, ‘neurotrophin signaling pathway’, ‘RNA degradation’, ‘GABAergic synapse’ and ‘ErbB signaling pathway’ (Figure 5B). Additionally, many of the downregulated KEGG terms (see Supplementary Table 3) also potentially show that the DG has more genes involved in mRNA production/degradation, alternative splicing and protein synthesis.

#### CA2xDG

The top results of KEGG pathways for upregulated genes in CA2 showed terms like: *‘pathways of neurodegeneration’, ‘thermogenesis’, ‘endocytosis’, ‘calcium signaling pathway’, ‘oxidative phosphorylation’, ‘carbon metabolism’, ‘glutamatergic synapse’, ‘synaptic vesicle cycle’, ‘GABAergic synapse’, ‘glycolysis/gluconeogenesis’,* and *‘citrate cycle (TCA cycle)’* (Figure 5C). On the other hand, top KEGG pathways for downregulated genes in CA2was: *‘ribosome’, ‘axon guidance’, ‘cAMP signaling pathway’, ‘signaling pathways regulating pluripotency of stem cells’, ‘spliceosome’, ‘neurotrophin signaling pathway’, ‘cell cycle’, ‘RNA degradation’,* and *‘TGF-beta signaling pathway’* (Figure 5D).

#### CA3xDG

Among the most enriched KEGG pathways for positively regulated genes in CA3 found: *‘pathways of neurodegeneration’, ‘endocytosis’, ‘thermogenesis’, ‘oxidative phosphorylation’, ‘calcium signaling pathway’, ‘carbon metabolism’, ‘dopaminergic synapse’, ‘synaptic vesicle cycle’, ‘pyruvate metabolism’,* and *‘citrate cycle (TCA cycle)’* (Figure 5E). Moreover, we did the KEGG analysis for downregulated genes in CA3 and found terms like: *‘MAPK signaling pathway’, ‘axon guidance’, ‘spliceosome’ ‘cAMP signaling pathway’,* ‘signaling pathways regulating pluripotency of stem cells’, *‘spliceosome’, ‘neurotrophin signaling pathway’, ‘cell cycle’, ‘RNA degradation’, ‘TGF-beta signaling pathway’,* and *‘mRNA surveillance pathway’* (Figure 5F).

## 4 DISCUSSION

In the present study we used the LCM technique coupled with RNA-seq to identify genes that are differentially expressed when comparing different subfields of the rat hippocampus. We aimed to separately analyze CA1, CA2, CA3 pyramidal layers, and DG granular layer to contrast the transcriptional profile of these neuronal populations. Previous studies have already described differences and similarities between pyramidal and granular cells (Cembrowski et al., 2016; Greene et al., 2009; Lein et al., 2004) but we are the first to employ RNA-seq for pairwise comparison of the rat hippocampus subfields and to quantify the differences in expression levels. Results obtained in this study provides data that may allow a deeper understanding of the rat hippocampus based on biological processes terms and quantitative transcriptional differences.

In order to identify potential hippocampal cell population markers, we searched for genes with an expression four times higher (>4 fold change) when comparing a subfield to all others. Notably, although some of our identified marker genes were previously described as markers in the literature, the majority of the discovered marker genes were novel for the rat hippocampus. In an example of potential marker genes, we have identified the gene *Pex5l* (peroxisomal biogenesis factor 5-like) for CA1, which is crucial to the establishment of a dendritic gradient of HCN1 channels and contributes to hyperpolarization in these neurons (Piskorowski et al., 2011). In another example, we also found *Rfx3* (regulatory factor X3) with higher expression in DG, which could directly regulate *Fgf1* (fibroblast growth factor 1) (Hsu et al., 2012) and contribute to DG neurogenesis (Ma et al., 2009). Therefore, our data considerably increased the total number of potential marker genes for each of the rat hippocampal subfields. Furthermore, our dataset may contain even more genes to be considered as potentially region markers if a less stringent criteria is used. We compared these genes against previously known markers from (Cembrowski et al., 2016) and found similarities and divergences between rat and mice hippocampus. For instance, our dataset has not identificated *Maob* (monoamine oxidase B) and *Scgn* (secretagogin, EF-hand calcium binding protein) as a CA2 marker gene or *Fybcd1* (fibrinogen C domain containing 1) as CA1 marker gene but, like (Cembrowski et al., 2016), we have identificated *Wfs1* (wolframin ER transmembrane glycoprotein) and *Fgf2* (fibroblast growth factor 2) as potential markers for CA1 and CA2 respectively. Intriguingly, recent studies have shown different protein expression patterns in the mouse and the rat hippocampus (Münster-Wandowski et al., 2017; Radic et al., 2017). However, since we used DESeq2 normalization and (Cembrowski et al., 2016) used FPKM normalization, it is not possible to rule out that part of the differences that we found are related to the normalization method rather than the animal model. Thus, our dataset recapitulated previous findings, but revealed unidentified genes that may directly demonstrate the differences between rat and mouse hippocampus.

We found that the comparison that has the largest number of DEGs is CA3xDG (Figure 1A). In addition, the PCA emphasizes the greatest variability between CA3 and DG in both PC1 and PC2 axes (Figure 1B). Interestingly, 3193 out of 7120 DEGs in CA3xDG overlap in CA2xDG and CA1xDG comparisons (Figure 1C), which shows how pyramidal and granule neurons have well-defined contrasting expression patterns. We also found that CA1xCA3 has the largest number of genes differentially expressed in the pyramidal comparison. On the other hand, CA2xCA3 shows 1847 DEGs, being more similar than any other comparisons. These subfield gene variations were already expected due to the role of anatomical differences, connections, firing properties (Kesner & Rolls, 2015; Mizuseki et al., 2012), and distinctive gene profiles on the hippocampus that each subfield has (Cembrowski et al., 2016). In summary, our data suggest that CA3 neuronal expression profiles are remarkably more distinct to CA1 and DG, however, CA3 shares more similarity to CA2.

Next, we focused on the ontology analysis to characterize the profile of all DEGs in the pyramidal comparisons. Our results indicate that the majority of significant DEGs are enriched for *‘ATP metabolic processes’* and *‘generation of precursor metabolites and energy’* for CA2 and CA3 subfields when compared to all other regions. These findings are consistent with other studies that point to more transcripts related to energetic processes, like glycolysis enzymes in the CA3 subfield (Datson et al., 2004; Greene et al., 2009). Our findings indicate that genes typically related to glycolysis, oxidative phosphorylation, gluconeogenesis, and lipid catabolic process are more abundant in CA2 and CA3 subfields. We also found that CA2 and CA3 neurons have a higher level of expression of genes responsible for the regulation of oxidative stress such as *Atox1* (antioxidant 1 copper chaperone), *Sod1/Sod2* (superoxide dismutase 1 and 2), *Aldh1a1* (aldehyde dehydrogenase 1 family member A1), and all thioredoxin reductases. Our results were similar to those of Yin et al. 2017, (Yin et al., 2017) which shows the greater reducing capability of CA3 than CA1 due to higher expression of thioredoxin reductases. We also hypothesized that the same may occur in the CA2 subfield since they share many similarities to the CA3 subfield. Interestingly, CA1 neurons are more sensitive than CA3 neurons to oxidative stress (Lana et al., 2020) and a low expression of oxidative stress regulators genes may be responsible for the lack of resistance of CA1 neurons to the damage caused by any inhibition of mitochondrial activity in these cells. In our study, CA1 pyramidal neurons have shown a lower expression of metabolism related genes when compared to other subfields. However, it has already been demonstrated that CA1 has a higher firing rate and energy demand than CA3 neurons (Mizuseki et al., 2012). A possible explanation for a higher expression of energy metabolism genes is due to the fact that CA2 and CA3 neurons are relatively larger than CA1 (Ishizuka et al., 1995). Furthermore, CA1 is vascularized by smaller and shorter vessels compared to the arteries supplying the CA3 subfield (“Vascular Patterns of the Rat Hippocampal Formation,” 1976), which might result in a higher oxygen supply for CA3 neurons. We also found that synaptic vesicle cycle genes, such as *Syp* (synaptophysin), *Vamp1* (vesicle-associated membrane protein), and *Syt11* (synaptotagmin 11) are more abundant in CA3 and CA2 and may correlate to intense energy metabolism and capability of greater synaptic vesicle release and/or recycling in these neurons (Pathak et al., 2015; Yuan et al., 2018). Here we report that CA2 and CA3 pyramidal neurons express more genes responsible for energetic processes, regulation of oxidative stress, and synaptic vesicle cycle.

Actin regulation is a crucial part of neural development for the growth of dendrites and axons. Our analysis indicates more transcripts related to *‘synapse organization’* and *‘actin filament organization’* in CA1. We found actin cytoskeleton signal transducers *Rnd1* and *Rnd3* (Rho GTPases 1 and 3) more abundant in CA1, implicating in a possible higher regulation of neuronal morphogenesis in this subfield (Luo, 2000). In addition, we also found important genes involved in neurite formation like *Map2* (microtubule-associated protein 2), *Twf2* (twinfilin actin-binding protein 2), and *Tpm1* (tropomyosin 1) (Gray et al., 2017; Leif Dehmelt, 2005; Yamada et al., 2007). Furthermore, we found that the transcription factor *Srf* (serum response factor), a classical regulator of several cytoskeletal genes, has a higher expression CA1 when compared to other subfields. *Srf* is critical to induce a gene expression pattern involved in the maintenance of long-term potentiation (LTP), since CA1 pyramidal neurons missing *Srf* exhibit an attenuation of both the early and late phases of LTP (Ramanan et al., 2005). Moreover, we found in CA1 an upregulation of *Myosin Vb,* a protein that has been established as a functional connection to recycling endosomes (Hammer & Sellers, 2011). Interestingly, recycling endosomes contribute to the regulation of AMPA receptors to the plasma membrane (Park et al., 2004) and in the present dataset the CA1 subfield has a higher expression of all four AMPA subunits(*Gria1, Gria2, Gria3, Gria4)* than CA2, CA3, and DG. Spine enlargement dependent on actin increases AMPA-receptor exposure in the membrane, further modulating LTP (Matsuzaki et al., 2004). Thus, the CA1 subfield expression of more actin regulatory genes may be related to LTP regulation and AMPA receptors.

CA2 has remarkable characteristics that differ from other CA subfields, like the lack of LTP and reduced synaptic plasticity (M. Zhao et al., 2007). However, in our study, CA2 was also enriched for *‘actin filament organization’* and *‘synapse organization’* genes in the CA2xCA3 comparisons. Therefore, the lack of synaptic plasticity in CA2 may not be explained by a lack of expression of genes associated with these functions. Compared to CA1 and CA3, CA2 neurons are known to be more resistant to trauma, seizure activity, and ischaemic insult, but the mechanisms responsible for such differences are not yet fully characterized (Dudek et al., 2016). Among the transcripts involved in cell survival and apoptosis, we identified that CA2 has a higher expression level of *ATP8A1* (ATPase phospholipid transporter 8A1), *TP73* (tumor protein p73), *Nes* (Nestin), and *CD74.* The lack of *ATP8A1* in hippocampal neurons is related to externalization of phosphatidylserine (Levano et al., 2012), which is associated with apoptosis (Fadok et al., 2000). Higher expression of *ATP8A1* in CA2 neurons may reduce phosphatidylserine externalization and subsequent programmed cell death, offering a mechanism of neuroprotection. Furthermore, one critical component of the development, maintenance, and survival of neurons is the *p73* pathway, which may also play a role in CA2, since it has a higher expression in this subfield. Deletion of *TP73* isoforms results in hippocampal dysgenesis and an impaired organization of CA1 and CA3 subfields (Yang et al., 2000). In addition, our data also show the upregulation of calcium-regulating proteins in CA2, such as *Casr* (calcium-sensing receptor), *Trpc3* (transient receptor potential cation channel C3), Cacng5 (calcium voltage-gated channel auxiliary subunit gamma 5), *Cacna2d3* (calcium voltagegated channel auxiliary subunit alpha2delta 3), and *RGS14* (regulator of G protein signaling 14), a classical modulator of calcium signaling and LTP suppression in this hippocampal subfield (Evans et al., 2018; Lee et al., 2010). Similar findings are represented in other studies and support the idea that the lack of LTP is related to CA2 neurons expressing a large number of calcium-regulating proteins (Simons et al., 2009). Therefore, these transcripts may present neuroprotective effects and play a role in intracellular signaling in this cell population.

A higher expression of protein synthesis genes in DG granule cells is well-known and was described in other studies (Datson et al., 2004; Greene et al., 2009). In our data, enriched BP terms for DG upregulated genes were mostly related to the *‘ribosome’, ‘RNA degradation’,* and *‘spliceosome’* and we found a large number of ribosomal proteins in DG, such as *Rpl3, Rpl4, Rpl5, Rpl7, Rps15, Rps16, Rps24* and *Rps28.* A higher expression level in the DG of genes involved in ribosomal biogenesis may indicate a robust ribosomal apparatus supporting dendrites’ growth and maintenance (Slomnicki et al., 2016). The DG subfield is capable of neurogenesis throughout life (Ninkovic et al., 2007) and protein synthesis may play a role in proliferation/differentiation and contribute to neuronal turnover in this region. In addition, we confirm the higher expression of pluripotency and neurogenesis molecular biomarkers like *Prox1* (prospero homeobox 1), *FoxO3* (forkhead box O3), *Calb1* (calbindin 1), and GEAP(glial fibrillary acidic protein) (Zhang & Jiao, 2015). We also found upregulation of neurotrophins like *BDNF* (brain-derived neurotrophic factor) and *NT-3* (neurotrophin-3), essential molecular mediators to stimulation of protein synthesis and synaptic plasticity in the nervous system (Aakalu et al., 2001). Furthermore, our transcriptomic analysis shows an abundance of alternative splicing factors in DG. Alternative splicing is an important mechanism that regulates the transcript isoforms and generates protein diversity in the cell (Su et al., 2018). We found an upregulation of *SRSF1* (serine and arginine rich splicing factor 1), *SRSF2* (serine and arginine rich splicing factor 2), *SRSF3* (serine and arginine rich splicing factor 3), *hnRNP* U (heterogeneous nuclear ribonucleoprotein U), *hnRNPa1* (heterogeneous nuclear ribonucleoprotein A1), and *Rbfox1* (RNA binding fox-1 homolog 1) that suggest a greater regulatory mechanism for mRNA isoforms. This study is consistent with previous findings that protein synthesis is increased in the granular neurons of DG, but sheds light that splicing factors are enhanced in DG as well.

The transcriptome data generated in the present study uncover the functional profiles and potential molecular mechanisms that underlie the distinction in CA1, CA2, CA3, and DG of rat hippocampal subfields. The gene ontologies enriched in this dataset reveal multiple biological processes such as actin regulation genes and AMPA receptors for CA1; several metabolic processes for CA2 and CA3 transcripts, and ribosome/spliceosome for DG. Many biological processes found enriched in the present dataset have been previously associated with specific characteristics of each hippocampal subfield, however here we present an extensive list of molecular components for each of these processes. Furthermore, the RNA-Seq approach allowed us to measure precisely the expression levels of transcripts from a large set of genes, revealing unique expressed genes for each subfield, contributing to a huge number of novel potential markers. The present findings contribute to a deeper understanding of the differences in the molecular machinery expressed by the rat hippocampal neuronal populations, further exploring underlying mechanisms responsible for each subfield specific functions.

## ACKNOWLEDGEMENTS

This work was supported by grants from Coordenação de Aperfeiçoamento de Pessoal de Nível Superior (CAPES) and Fundação de Amparo à Pesquisa do Estado de São Paulo, SP, Brazil (FAPESP; grant number 2013/07559-3)

## AUTHOR CONTRIBUTIONS

**João P. D. Machado:** Conceptualization, Methodology, Investigation, Visualization, Formal analysis, Writing—Original Draft. **Alexandre H. B. Matos and Iscia Lopes-Cendes:** Conceptualization and Methodology, Writing—Original Draft. **Maria C.P. Athie:** Conceptualization, Methodology, Formal analysis, Writing—Original Draft. **André. S. Vieira:** Conceptualization, Methodology, Project administration, Supervision, Writing-Original Draft.

## DATA AVAILABILITY STATEMENT

The data that support the findings of this study are available from the corresponding author upon reasonable request.

## Supplementary Information

**Supplementary table 1** – List of genes differentially expressed in all subfield comparisons.

**Supplementary table 2** – List of possible marker genes with four times more expression (>2 log2FoldChange) in only one subfield.

**Supplementary table 3** – List of biological processes from GO analysis significantly enriched based on all, up or down-regulated genes in CA1, CA2, CA3, and DG subfields.

**Supplementary table 4** – List of KEGG pathways significantly enriched based on all, up or down-regulated genes in CA1, CA2, CA3, and DG subfields.

